# APC loss promotes intestinal transformation through induction of bistable stem cell states

**DOI:** 10.1101/2025.03.06.641686

**Authors:** N Li, RL Myers, Q Zhu, Y Tian, K Durning, X Wang, NA Leu, JH Rhoades, D Bankler-Jukes, KE Monaghan, S Adams-Tzivelekidis, O Pomp, O Pellon-Cardenas, MA Blanco, MP Verzi, PS Klein, CJ Lengner

## Abstract

Colorectal cancer (CRC) is a leading cause of cancer deaths, predominantly initiated by genetic inactivation of the APC tumor suppressor. Current dogma holds that APC inactivation promotes tumorigenesis via hyperactivation of the WNT pathway transcriptional effector, β-CATENIN. Although β-CATENIN activation is required for tumor initiation, activating mutations in β-CATENIN are infrequent. Here, we ask what underlies the selective pressure for APC inactivation by comparing the oncogenic effects of APC loss to β-CATENIN hyperactivation. We find that APC loss activates a β-CATENIN-independent fetal intestinal transcriptional program driven by dysregulation of GSK-3 activity upon the LIM-domain protein AJUBA, a positive regulator of YAP. This results in AJUBA stabilization and downstream transcriptional activation of the YAP-driven fetal intestinal gene expression program. We find that β-CATENIN- and YAP-driven transcriptional states are mutually exclusive, existing in an interchangeable bistable balance among APC-null cells. This results in more robust tumor initiation and metastatic progression downstream of APC loss relative to β-CATENIN activation. Taken together, our findings explain the preferential selection for APC inactivation in CRC development and illuminate how β-CATENIN- and YAP-driven gene expression programs coexist to promote tumorigenesis.

## INTRODUCTION

Colorectal cancer (CRC) is among the most diagnosed cancers and is a leading cause of cancer-related deaths, with both incidence and mortality increasing among younger individuals and in developing countries. Genetic inactivation of the APC tumor suppressor is the most frequent genetic lesion in CRC (mutated in over 80% of microsatellite-stable CRC cases), where it is believed to be the initiating event^1^. APC has a well-established role as a negative regulator of β-CATENIN, the transcriptional effector of the canonical WNT signaling pathway. In this context, APC coordinates the activity of the cytoplasmic destruction complex- a multiprotein complex that also includes AXIN, GSK-3, and CK1α^2^. In the intestinal epithelium, in the absence of Wnt ligand-receptor interaction, GSK-3, and CK1α continually phosphorylate β-CATENIN within the destruction complex, priming it for subsequent ubiquitylation by the ubiquitin ligase β-TRCP, ultimately resulting in its proteasomal degradation. Upon WNT ligand engagement with FZD receptors, APC, GSK-3, and the destruction complex disengage β-CATENIN, allowing it to accumulate and enter the nucleus where it functions as a transcription factor in conjunction with TCF/LEF cofactors, promoting the expression of pro-mitotic and intestinal stem cell-associated genes including *CCND1*, *ASCL2, MYC*, and *LGR5*^3^.

APC mutations driving CRC (C-terminal truncations and large genic deletions) result in the destruction complex disengaging β-CATENIN, causing pathogenic β-CATENIN accumulation and target gene activation^4,5^. Interestingly, Samowitz et al.^6^ observed that β-CATENIN activating mutations are more frequent in small adenomas than large adenomas, and are infrequently associated with invasive cancer, relative to APC inactivating mutations. These observations led them to propose, some 25 years ago, that APC inactivating mutations are not functionally equivalent to β-CATENIN activating mutations, and that APC loss drives other pathways to cause malignant disease.

APC loss in intestinal stem cells promotes hyperplasia and adenoma formation, and these phenotypes require downstream MYC activity^7^. β-CATENIN and MYC activation are well-established, requisite oncogenic consequences of APC loss; however numerous lines of evidence support β-CATENIN-independent functions of APC. Such β-CATENIN-independent activities include microtubule organization^8^, such that APC inactivation perturbs epithelial polarity and mitotic spindle orientation^9,10^. They also include RNA binding functions associated with decreased translation, although this function has not been investigated in the context of the intestinal epithelium^11^. APC also promotes GSK-3 activity toward targets beyond β-CATENIN^12^. This includes TSC2, a negative regulator of mTORC1, resulting in mTORC1 activation upon APC loss^13,14^. APC and GSK-3 also suppress YAP (the transcriptional effector of Hippo signaling) by enhancing the activity of Large Tumor Suppressor Kinase 1 (LATS1), although the direct target of GSK-3 in this regulation has not been identified^15^. Ultimately, a clear demonstration of the β-CATENIN-independence of these APC functions in epithelia has been challenging due to an absolute requirement for β-CATENIN in the adherens junctions of epithelia. This precludes the use of genetic inactivation of *CTNNB1* (the gene encoding β-CATENIN) to dissect the contribution of β-CATENIN to the oncogenic phenotypes downstream of APC inactivation.

Here, we set out to define β-CATENIN-independent consequences of APC loss that contribute to CRC initiation and progression. We demonstrate that APC loss drives a gene expression program normally associated with fetal intestinal stem cell (ISC) activity (reviewed in^16^). APC-null cells enriched for this program exhibit increased stem cell self-renewal in organoid assays relative to cells with oncogenic β-CATENIN gain-of-function, and APC-null cells exhibit enhanced tumor initiation and metastasis capacity *in vivo*. We provide evidence that activation of fetal gene expression downstream of APC loss is independent of β-CATENIN through the generation of a drug-inducible mouse model specifically inhibiting β-CATENIN transcriptional activity. We describe how APC loss impairs GSK-3-mediated phosphorylation of the LIM-domain protein AJUBA, resulting in its stabilization. AJUBA is a negative regulator of LATS1, which phosphorylates YAP, promoting its cytosolic localization and targeting it for degradation. Thus, GSK-3 phosphorylation of AJUBA provides a molecular link between APC and LATS1-mediated YAP degradation: In the absence of APC, GSK-3 cannot phosphorylate AJUBA, which accumulates and suppresses LATS1, ultimately resulting in YAP stabilization, nuclear localization, and activation of fetal gene expression independent of β-CATENIN.

We find that in APC-null epithelia, the β-CATENIN-driven canonical adult intestinal gene expression program and the YAP-driven fetal gene expression program are anticorrelated and nearly mutually exclusive. However, these programs co-exist at the population level to create a bistable balance in stem cell states, with cells readily switching states, providing insight into how plasticity between these two programs is governed by APC. Interestingly, in human CRC, we observe that expression of the fetal intestinal signature correlates with poor outcomes only in APC-mutant human CRC, but not in APC wildtype cancers.

Much attention has recently focused on plasticity of stem cell states in colorectal cancer^16–19^ and the role of YAP in promoting colon cancer^16,20–23^. More recently, the importance of YAP in promoting fetal intestinal gene expression programs and intestinal regeneration has become increasingly clear^24–27^. Our findings describe a novel oncogenic mechanism activated upon APC loss in a β-CATENIN-independent manner resulting in bistable epithelial stem cell states and offer a molecular explanation for the longstanding clinical observation that human colorectal cancers have a proclivity for inactivating APC rather than activating mutations in other components of the canonical WNT-β-CATENIN pathway.

## RESULTS

The *APC* gene is well-established as the most frequently mutated gene in colorectal adenocarcinoma. We initially examined publicly available cancer datasets^1^ for the mutation frequencies of *APC* and *CTNNB1* (encoding β-CATENIN) and found, as expected, a clear selection for *APC* mutation relative to *CTNNB1* (**Fig. 1A**). Interestingly, this pattern was restricted to CRC (**Fig. 1A**), suggesting a unique sensitivity of the colonic epithelium to *APC* loss, relative to other epithelial tissues-of-origin in carcinoma. We therefore sought to directly compare the effects of APC loss-of-function (LOF) to β-CATENIN gain-of-function (GOF) in mouse models. We selected an *Apc* floxed allele (*Apc^flox^*) in which the penultimate exon is deleted upon Cre induction-a widely employed model for CRC initiation^28^. For β-CATENIN GOF, we employ a *Ctnnb1* allele in which exon 3 is deleted upon Cre induction (*Ctnnb1^ex3flox^*)^29^. This model, also widely used, ablates the amino acids that are phosphorylation targets of the destruction complex responsible for β-CATENIN proteasomal degradation, and thus Cre-mediated deletion of this allele activates β-CATENIN similar to the mutations observed (albeit infrequently) in human CRC. We confirmed key findings using a targeted, doxycline-inducible, constitutively active β-CATENIN allele^30^.

**Figure 1.**
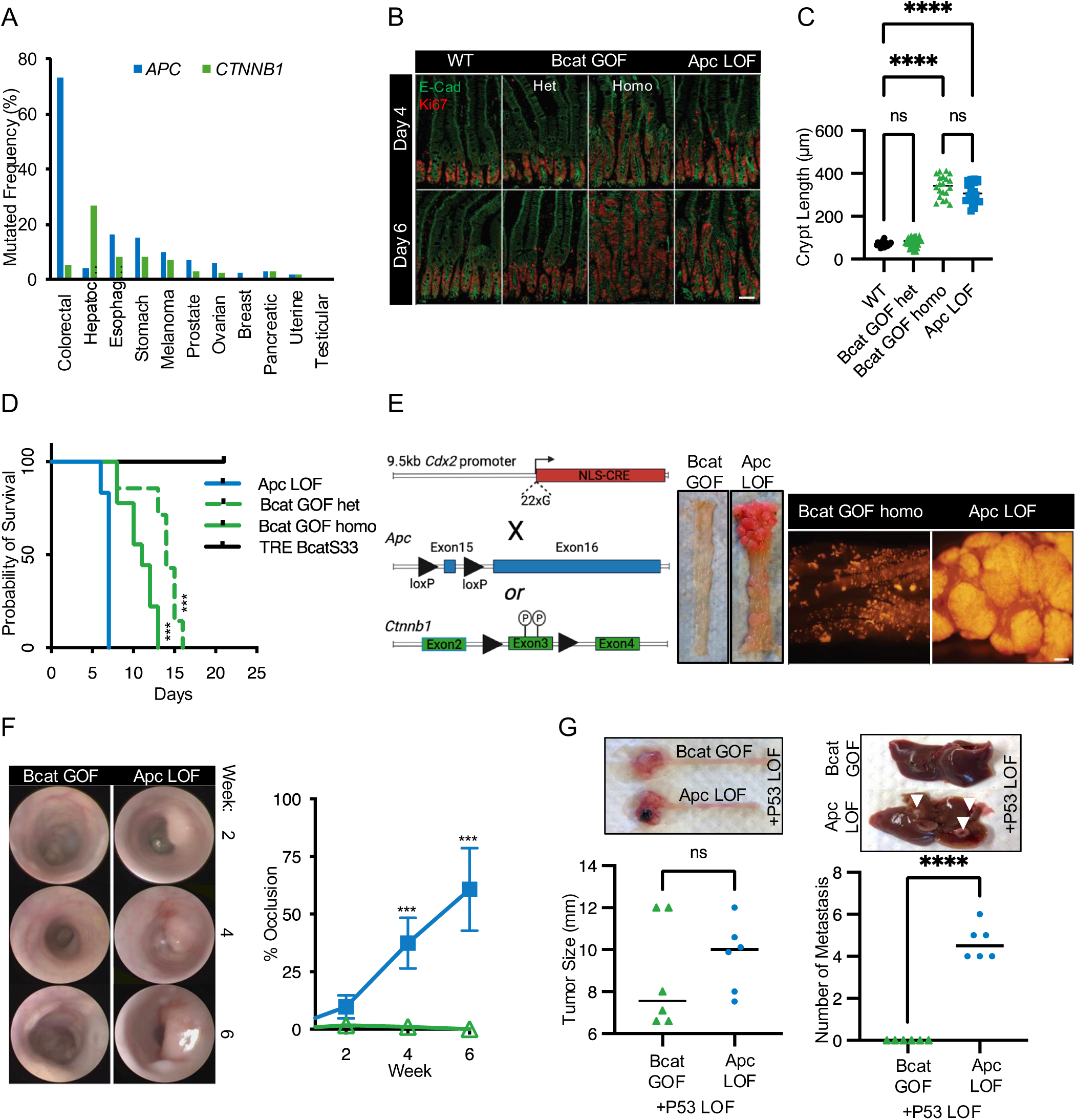
Comparison of intestinal epithelial transformation downstream of APC inactivation versus β-CATENIN activation. **A.** Mutation frequency analysis for *APC* and *CTNNB1* across 11 major human cancer types from TCGA data. **B.** Acute inactivation of APC or activation of β-CATENIN in response to Villin-CreER induction in the adult mouse jejunum at indicated timepoints, stained for epithelial E-CADHERIN and KI67. Scale bar, 50 µm. **C.** Quantification of expanding crypt length on day 4 after CreER induction. ****P < 0.0001, ns: not significant. **D.** Kaplan–Meier plot of *Apc^flox/flox^* (APC LOF), *Ctnnb1^ex3flox/ex3flox^* (Bcat GOF homo), and *Ctnnb1^ex3flox/wt^*(Bcat GOF het) mouse models after CreER activation, along with *TRE-β-CATENIN-S33* mice after doxycycline induction. Statistical significance (P < 0.001) is indicated for both Bcat GOF homozygous and heterozygous groups compared to Apc LOF mice. **E.** Schematic representation of genetic constructs and strategies used to inactivate APC or activate β-CATENIN using the *Cdx2-G22-Cre* transgene. Middle panel: Macroscopic appearance of colon of indicated genotype. Right panel: Fluorescent imaging of tdTomato from the *R26^lox-stop-lox-tdTomato^* allele in these mice, as a proxy for Cre activation, in whole mount colon tissue. Scale bar: 100 µm. **F.** Left panel: Representative endoscopic images of colonic lumen in Apc LOF and Bcat GOF mice at 2-, 4-, and 6-weeks post-injection, quantified at right. ***P < 0.001. **G.** Top panels: Gross examination of primary colon tumors (left) and macroscopic metastatic liver lesions (right, arrowheads) in Bcat GOF and Apc LOF mice with P53 LOF. Lower panel: Quantification of primary tumor size (left) and metastatic lesion frequency in Bcat GOF + P53 LOF and Apc LOF + P53 LOF mice. ****P < 0.0001.

We initially examined the immediate and direct consequence of APC LOF vs. β-CATENIN GOF by crossing *Apc^flox^* and *Ctnnb1^ex3flox^* to a pan-epithelial, tamoxifen-inducible Cre transgenic allele, *Villin-CreER*^31^. Consistent with prior studies^28,32^, APC LOF in *Apc^flox/flox^*::*Villin-CreER* mice resulted in rapid crypt hyperplasia and mortality within one week of tamoxifen induction (**Fig. 1B-D**). *Ctnnb1^ex3flox/wt^*::*Villin-CreER* mice exhibited minimal hyperplasia one-week post-tam induction, and did not exhibit mortality until at least two weeks post-tam induction. While homozygous *Ctnnb1* mutations are exceedingly rare, constitutively active alleles in *Ctnnb1^ex3flox/ex3flox^* mice were required to accelerate hyperplasia and mortality approaching that seen upon APC LOF (**Fig. 1B-D**). Interestingly, an orthogonal mouse model of constitutive β-CATENIN activation, a doxycycline-inducible, non-degradable β-CATENIN S33 mutant transgene (*TRE-BcatS33*)^30^ did not exhibit morbidity/mortality across the time course analyzed, despite evident crypt hyperplasia (**Fig. 1D** and **S1A**). We therefore employ *Apc^flox/flox^*::*Villin-CreER* and *Ctnnb1^ex3flox/ex3flox^*::*Villin-CreER* for most subsequent experiments and refer to them as APC LOF and β-CATENIN GOF. Both APC LOF and β-CATENIN GOF mice exhibited the established Paneth cell mislocalization known to be controlled by β-CATENIN^33^. Interestingly, APC LOF decreased numbers of Paneth cells in contrast to β-CATENIN GOF, and both models exhibited decreased enteroendocrine cell frequency (**Fig. S1B-E**). β-CATENIN GOF exhibited increased combined histopathological scores relative to APC LOF, despite the decreased mortality of APC LOF (**Fig. S1F**).

These studies suggest more pronounced effects of APC inactivation relative to β-CATENIN activation but preclude longitudinal analysis of tumorigenesis due to hyperplastic field effects related to broad Cre-mediated recombination and rapid mortality due to dehydration. We therefore turned to a clonal deletion model to compare the effects of APC LOF vs. β-CATENIN GOF on tumor formation. Different floxed alleles recombine with drastically different frequencies when using tamoxifen inducible Cre drivers^34^, precluding the direct quantification of tumor initiation downstream of APC LOF vs. β-CATENIN GOF using limiting/clonal CreER induction. We therefore employed a stochastic, constitutive Cre allele to compare tumor formation efficiencies between models. The *CDX2P9.5G22Cre* transgene^35^ employs a 9.5kb fragment of the *Cdx2* promoter, broadly expressed in the intestinal epithelium with increasing activity distally (colon), separated from a constitutive Cre coding sequence by a repeat sequence of 22 guanines (**Fig. 1E**). This places Cre out of frame until a stochastic S-phase error occurs, adding or removing guanines and thus occasionally placing Cre in frame. Because the Cre is constitutively expressed once in frame, this model will recombine different floxed alleles with analogous efficiency in contrast to limiting tamoxifen induction with CreER alleles. We also bred a *R26^lox-stop-lox-tdTomato^* allele^36^ into mice with either APC LOF or β-CATENIN GOF to monitor Cre activation upon the *CDX2P9.5G22Cre* transgene slipping into frame.

Remarkably, no *Apc^flox/flox^*::*CDX2P9.5G22Cre* mice survived at birth, and some *Apc^flox/wt^*::*CDX2P9.5G22Cre* mice exhibited visible tumors (**Fig. 1E** and **S1G**). This is in stark contrast with *Ctnnb1^ex3flox/ex3flox^*::*CDX2P9.5G22Cre* mice, which were born viable at near-Mendelian ratios but exhibited no tumors despite abundant Cre-mediated recombination events. Similarly, *Ctnnb1^ex3flox/wt^::CDX2P9.5G22Cre* mice were born at Mendelian ratios and exhibited no tumors (**Fig. 1E** and **S1G**). This finding confirms that APC loss has more potent tumor-promoting effects relative to constitutive β-CATENIN activation. We sought to confirm this observation with additional, orthogonal approaches. Both subcutaneous (**Fig. S1H**) and endoscope-guided orthotopic implantation (**Fig. 1F and S1I**) of APC LOF organoids into the distal colonic mucosa of syngeneic C57Bl/6 mice revealed robust adenoma growth, while implantation of β-CATENIN GOF (homozygous) organoids revealed little to no ability to support tumor formation.

All the models employed thus far investigate tumor initiation/adenoma formation. To compare the effects of APC LOF or β-CATENIN GOF on adenocarcinoma progression, we inactivated *Trp53*, encoding the P53 tumor suppressor (the second-most mutated gene in CRC often co-occurring with *APC*) using CRISPR/Cas9 in the adenomatous organoids. Orthotopic implantation of APC LOF::P53 LOF or β-CATENIN GOF (homozygous)::P53 LOF tumor organoids into syngeneic colonic mucosa drove formation of primary tumors in both genotypes (**Fig. 1G)**. Remarkably, APC LOF tumors exhibited robust metastatic spread to the liver, in contrast to those with homozygous β-CATENIN GOF in which no metastases were evident (**Fig. 1G**). These findings show that APC loss is more potent in promoting tumor initiation and progression relative to constitutive β-CATENIN activation, providing an explanation for the proclivity for APC inactivation in human colorectal cancers.

To understand the basis for the observed tumor phenotypes, we generated *Apc^flox/flox^*::*Villin-CreER* (APC LOF) and *Ctnnb1^ex3flox/ex3flox^*::*Villin-CreER* (β-CATENIN GOF) organoids and induced recombination *in vitro*. Surprisingly, while β-CATENIN GOF organoids proliferated and cultures were stably maintained, they exhibited drastically reduced clonal organoid formation efficiency (often used as a proxy for tumor initiation capacity/stem cell activity) relative to APC LOF organoids, resulting in dramatic differences in cumulative growth over serial passage (**Fig. 2 A-C and S2A**). Organoids of both genotypes contained differentiated cell types; however, cells within the APC LOF organoids exhibited decreased adhesion, with more fragmentation during passage and processing (**Fig. 2C**), which may relate to their increased metastatic proclivity described above. The observed differences in organoid growth might be explained by differences in cell cycling rates between genotypes, yet both S-phase progression and Ki67 reactivity revealed no differences in cycling between β-CATENIN GOF and APC LOF cultures (**Fig. 2E and S2B, C**).

**Figure 2.**
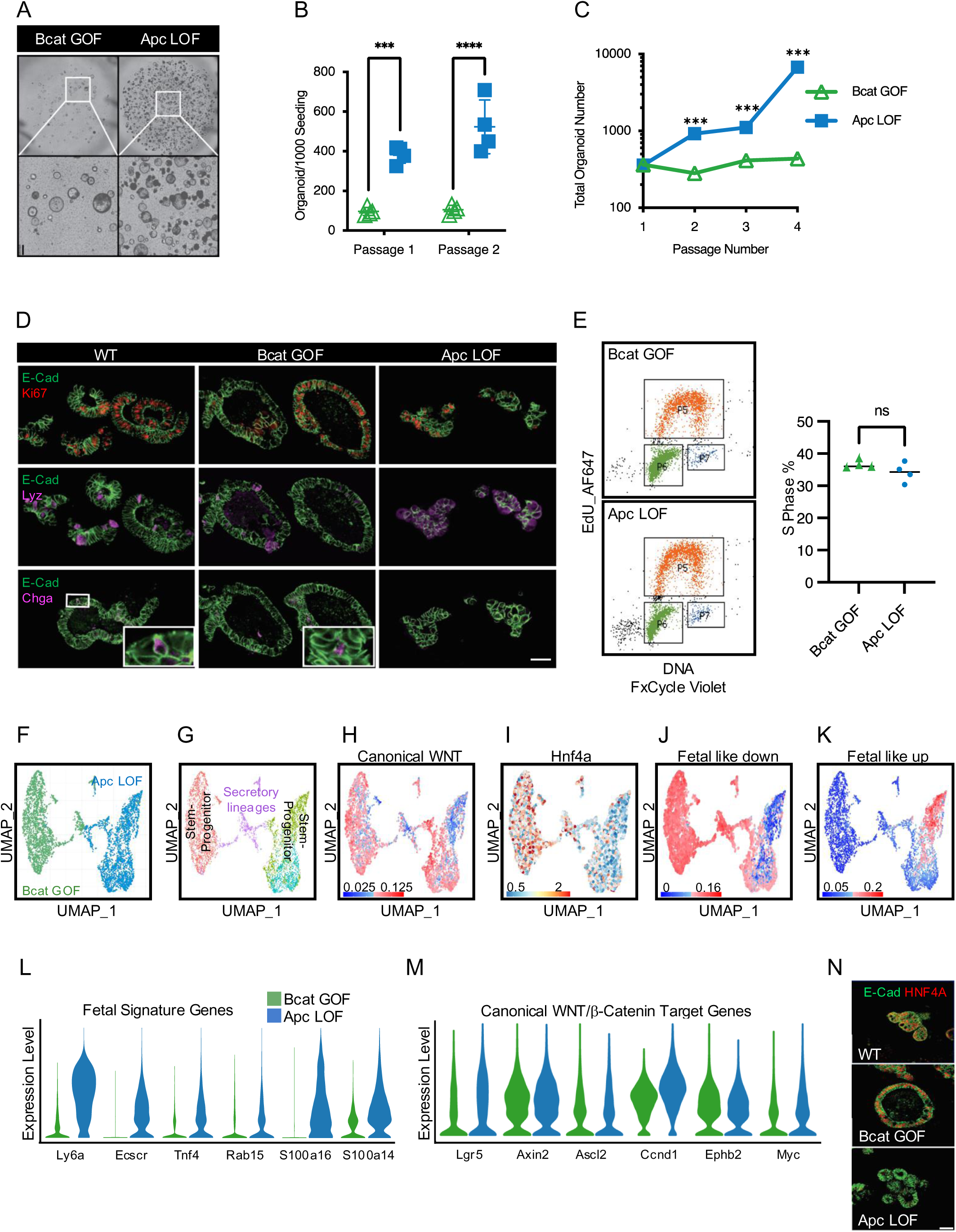
*Apc* inactivating mutations, but not *Ctnnb1* activating mutations, promote stem cell phenotypes. **A**. Organoid cultures derived from freshly isolated crypts 5 days after CreER-mediated recombination of target alleles. Scale bar, 50 µm. **B.** Clonal (single cell) organoid seeding efficiency in continuously passaged cultures. **C.** Cumulative organoid numbers after 4 passages in Apc LOF vs. Bcat GOF cultures. **D.** Staining for differentiation (Lyz and Chga) and proliferation (Ki67) markers in organoid cultures. Scale bar, 50 µm. **E.** Left panel: Analysis of S-phase progression in organoid cultures via EdU incorporation, quantified at right. **F.** Uniform Manifold Projections (UMAP) of single cell transcriptomes in Apc LOF vs. Bcat GOF organoid cultures. **G-K**. UMAP plots as in F showing cell type distribution (G), enrichment for the canonical WNT gene set (H), and Hnf4a gene (I), and gene sets derived from fetal versus adult organoid transcriptome profiles enriched in adult (fetal like down) (J) or fetal (fetal like up) (K) organoids. **L, M.** Violin plots showing expression of hallmark fetal signature genes (L) and canonical Wnt/β-CATENIN target genes (M). **N.** Immunofluorescence staining of organoids of indicated genotype for HNF4A and E-Cadherin. Scale bar= 50 µm. ****P < 0.0001, ***P < 0.001.

We next investigated the molecular underpinnings of the observed phenotypes through single cell transcriptomic analysis of β-CATENIN GOF and APC LOF organoid cultures. Uniform Manifold Approximation and Projection (UMAP) revealed that the major differences in transcript abundance between genotypes is driven by the stem/progenitor/absorptive cells, while cells of the secretory lineages were more transcriptionally similar across groups (**Fig. 2F, G**). Gene set enrichment analysis^37^ of these data revealed several things. First, levels of canonical WNT/β-CATENIN target transcripts were identical across genotypes, indicating that the phenotypic differences are not due to differing levels of β-CATENIN transcriptional activity^30^ (**Fig. 2H, M**). Next, APC loss was correlated with dramatic decreases in *Hnf4a*/ HNF4A RNA and protein, compared to β-CATENIN GOF (**Fig. 2I, N**). HNF4 enforces cell identity and differentiation (reviewed in ^38^) and has been implicated as a colorectal cancer tumor suppressor^38,39^. We confirmed HNF4 suppression *in vivo* in APC LOF adenomas relative to β-CATENIN GOF-driven tumors (**Fig. S2D**), and found that this was maintained in metastatic liver lesions in APC::P53 mutant tumors (**Fig. S2E**). HNF4 is associated with epithelial cell fate specification and differentiation, as is CDX2, and target gene sets for both transcriptional regulators (defined as genes with ChIPseq binding sites in mouse small intestine and downregulated in the knockout of the corresponding transcription factor^40,41^) were efficiently suppressed upon APC LOF relative to β-CATENIN GOF (**Fig. S2F**). This was further evidenced by suppression of individual genes related to differentiation including tight junctional gene *Cldn15*, the transcriptional regulator of intestinal identity *Cdx1*, and genes involved in digestion/absorption *Mttp* and *Aldob* (**Fig. S3A**). Thus, suppressed absorptive lineage differentiation may contribute to increased stem cell activity observed by clonal organoid formation assay and increased tumorigenesis observed upon APC LOF relative to β-CATENIN GOF (**Fig. 2A-C**).

Finally, and most strikingly, APC loss drove a dramatic increase in the expression of fetal intestinal genes (**Fig. 2J-L and S3B-F**). In contrast, fetal transcripts were barely detectable in β-CATENIN GOF or wildtype cultures (**Fig. 2J-L and S3B-F**). We also confirmed these effects were not unique to the *Ctnnb1^ex3flox^* model, as the inducible *TRE-BcatS33*^30^ behaved similarly to *Ctnnb1^ex3flox^* (**Fig. S3F**).

While these data demonstrate that constitutive β-CATENIN activation is insufficient to phenocopy APC loss - particularly the striking activation of fetal gene expression programs - β-CATENIN activity may still be necessary to induce fetal gene expression upon APC LOF. To formally test this, we targeted a single-copy, doxycycline (Dox)-inducible, dominant negative TCF4^42^ (dnTCF4) transgene into safe-haven chromatin in mouse embryonic stem (ES) cells which harbor a *R26-M2rtTA* allele-a system that enables robust Dox-inducible expression in the intestinal epithelium^43–45^ (**Fig. 3A**). We verified Dox-inducible dnTFC4 expression (**Fig. S4A**) and derived the novel *Col1a^TetOdnTcf4^::R26^m2rtTA^* mouse strain, from these ES cells, hereafter referred to as *TRE-dnTcf4* (Tetracycline-Responsive Element-dominant negative Tcf4). We crossed the *TRE-dnTcf4* alleles into *Apc^flox/flox^*::*Villin-CreER* mice, derived organoids, inactivated APC, then activated *TRE-dnTcf4* with 48hrs Dox treatment, followed by single cell transcriptome sequencing. As expected, induction of dnTcf4 efficiently silenced canonical WNT/β-CATENIN target genes (**Fig. 3B-D**). Strikingly, the induction of fetal gene expression programs downstream of APC loss was unaffected by inhibition of β-CATENIN transcriptional activity (**Fig. 3B-D** and **S4B**). Despite this robust induction of fetal gene expression, long-term survival of APC GOF cultures remained dependent on β-CATENIN transcriptional activity, consistent with the literature describing a requirement for β-CATENIN transcriptional activity, and activity of its target gene *Myc* for tumorigenesis downstream of APC loss^7^ (**Fig. S4C**).

**Figure 3.**
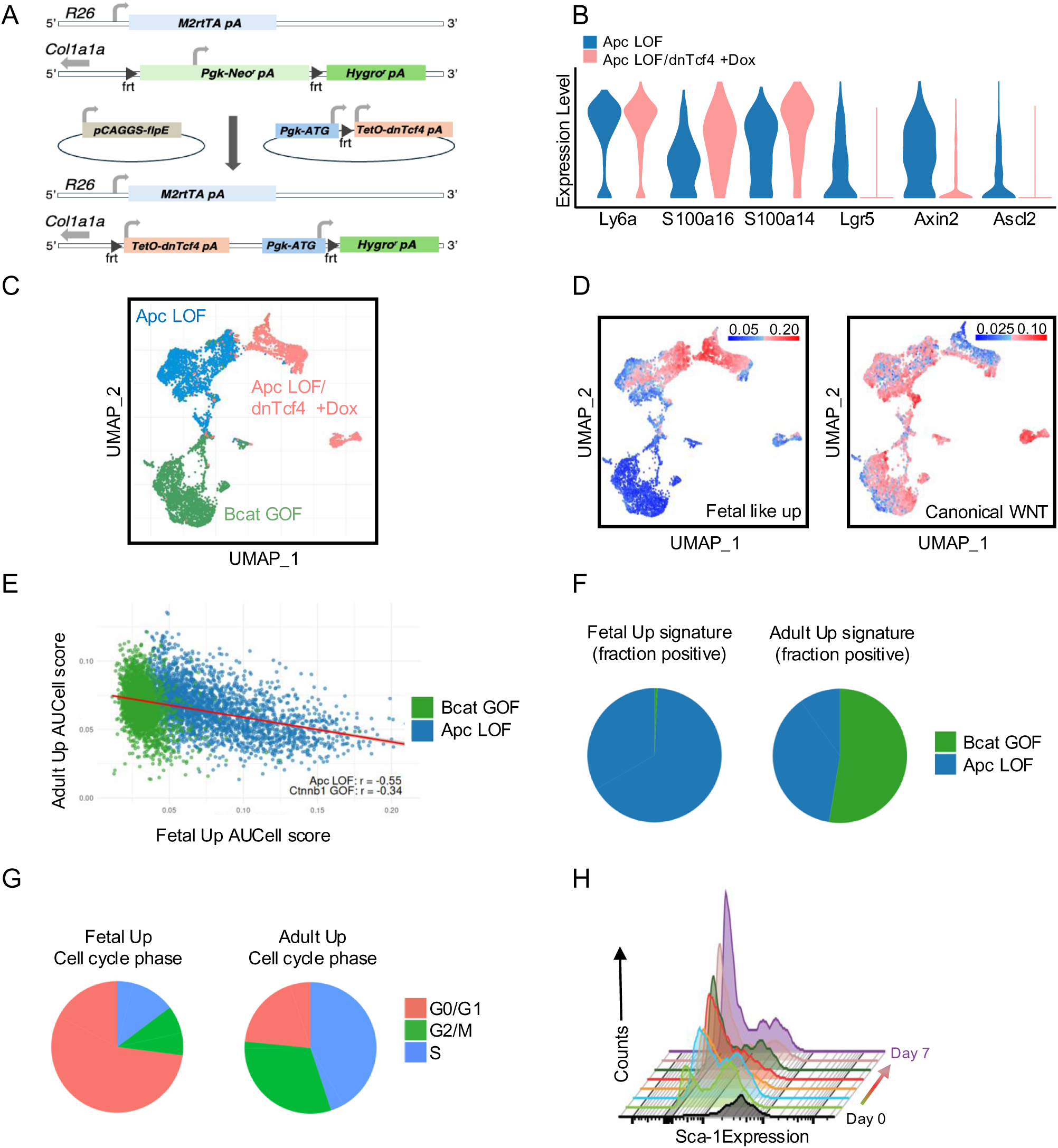
APC inactivation induces a fetal stem cell state independent of β-CATENIN, resulting in bistable fetal-adult stem cell populations. **A.** Schematic of the single-copy, targeted transgenic alleles for driving doxycycline-inducible expression of dominant-negative TCF4 (dnTcf4). **B.** Violin plots from single cell transcriptome profiles of Apc LOF:: TetO-dnTcf4 organoids after doxycycline administration, examining hallmark fetal genes (*Ly6a, S100a16, S100a14*) and adult, canonical WNT/β-CATENIN target genes (*Lgr5, Axin2, Ascl2*). **C.** UMAP plot showing Bcat GOF, Apc LOF, and Apc LOF:: TetO-dnTcf4+dox cells. **D.** UMAPs as in (C) showing enrichment for fetal intestinal and canonical WNT gene signatures. **E.** Scatter plot illustrating the relationship between the AUCell scores for fetal signature and adult signature genes. **F.** Pie graphs indicated the fraction of cells enriched for fetal or adult intestinal stem cell signatures based on genotype Bcat GOF vs. Apc LOF. **G.** Cell cycle phase distribution for cells expressing the fetal vs. adult gene expression signatures. **H.** Histograms derived from a flow cytometric timecourse in which Sca1+ cells are sorted from Apc LOF cultures and replated, monitoring Sca1 expression across one week of organoid growth.

The fetal intestinal gene expression program was originally defined by comparative transcriptomics between intestinal organoids derived from adult and fetal epithelium^46^. This program is essentially silent in the healthy adult intestine but becomes strongly induced in response to a variety of injuries, when the canonical WNT program becomes suppressed^26^. In the context of regenerating healthy tissue, this fetal program is driven, at least in part, by macrophage-derived TGFβ, ultimately resulting in the YAP transcription factor driving fetal gene expression^25,27^. Further studies revealed reactivation of this program in colorectal adenocarcinomas, with TGFβ again being implicated as an upstream cytokine driver and YAP, along with SOX9 executing the downstream transcriptional activation of fetal gene expression^16–19,23,47^. Our findings indicate that APC loss alone is sufficient to induce fetal-like reversion of the adult epithelium, and that this program is cell-autonomous, independent of cues from the tumor microenvironment, and independent of β-CATENIN activity.

While APC LOF is capable of concomitantly activating both the fetal and adult (e.g., β-CATENIN-driven) stem cell programs, a closer examination of our single cell transcriptomic data reveals these programs to be largely mutually exclusive among individual cells within the APC LOF population (**Fig. 2H**, **J**, **K**, and **Fig. 3E**). Cells expressing the fetal program are essentially exclusive to APC LOF cultures, while the cells expressing the adult program can be found similarly in both APC LOF and β-CATENIN GOF cultures (**Fig. 3F**). Interestingly, while both cultures proliferate at analogous rates overall (**Fig. 2E**), the fetal program was predominantly observed in cells in the G1 phase, while the adult program was found in cells more evenly distributed across the cycle, with a bias toward S-phase (**Fig. 3G**). These findings suggest that a cell’s residence in these differing states may be highly plastic downstream of APC inactivation. To test this, we separated cells in the fetal state based on their cell surface expression of Sca1 (encoded by the hallmark fetal signature gene *Ly6a,* **Fig. 2L**), replated Sca1^High^ and Sca1^-^ cells, and monitored distribution of cell states after culture. Baseline APC LOF cultures have a roughly equal distribution of cells in the adult and fetal states, and within one week of culture, fetal Sca1^High^ cells reestablished this equilibrium, giving rise to Sca1^-^ cells (**Fig. 3H and S4D**). Similarly, purified Sca1^-^ cells efficiently repopulated the Sca1^High^ population (**Fig. S4D**).

YAP transcriptional activity is linked to metastasis^48^, consistent with our findings in Fig. 1G, as well as in resistance to radiation or DNA damaging therapy^48,49^. We thus asked whether the fetal/ Sca1^High^ cells exhibited differential radioresistance relative to the adult/Sca1^-^ population in APC LOF cultures. While high doses (14 Gy) of ionizing radiation killed all cells, at 8 Gy, Sca1^High^ cells exhibited significantly increased survival relative to their Sca1^-^ counterparts (**Fig. S4E**).

Taken together, our findings indicate that APC inactivation activates the fetal gene expression program independently of β-CATENIN activity, creating epithelial populations that exist in a bistable stem cell state, fluctuating between fetal and adult intestinal stem cell identities. We also examined how expression of the fetal gene signature predicts survival in APC mutant and APC wildtype patients (the latter having other activating WNT mutations nearly universally^1^). Interestingly, fetal gene expression was not predictive of survival in APC wildtype patients, but becomes strongly correlated with poor survival when APC is mutated (**Fig. S4F**), indicating how this gene expression program contributes to tumor progression in CRC.

The fetal gene expression program is driven by YAP transcriptional activity in regenerative epithelium post-injury (reviewed in ^16^). We indeed observed stabilization of YAP protein and activation of the intestinal oncogene and YAP transcriptional target ZYMND8^50^, along with the expected suppression of HNF4A, in APC LOF organoids relative to WT or β-CATENIN GOF cultures (**Fig. 4A**). This was further confirmed by nuclear localization of YAP (**Fig. 4B**). We reasoned that YAP activation may arise through reduced GSK-3 activity due to APC loss. APC potentiates GSK-3 activity on numerous substrates beyond β-CATENIN^12,51^, including TSC2, a negative regulator of mTORC1, and thus mTORC1 is activated upon APC loss^14,13^. Indeed, we observed robust mTORC1 activity in APC LOF organoids relative to WT or β-CATENIN GOF cultures (**Fig. 4A**), confirming published findings.

**Figure 4.**
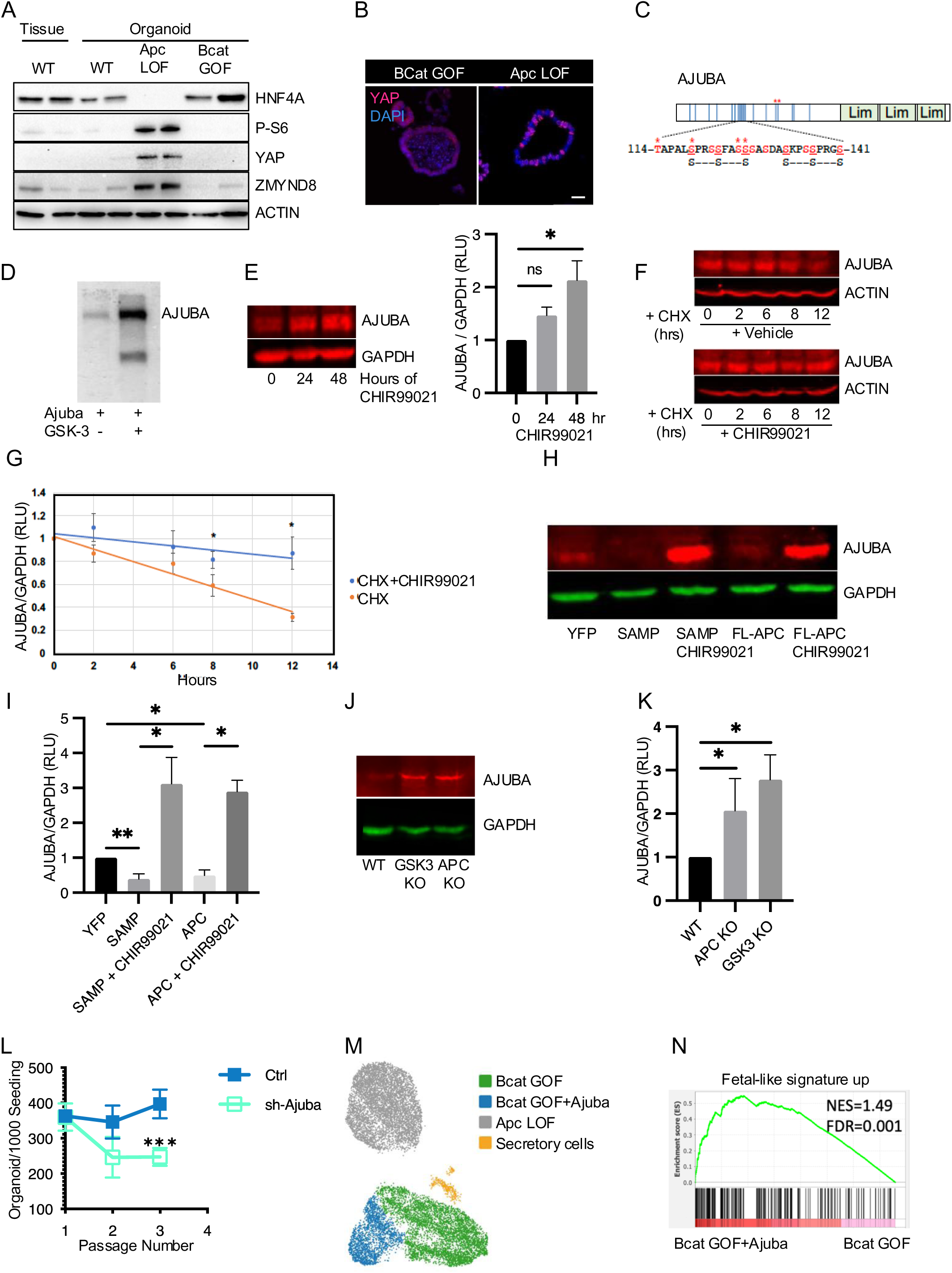
APC inactivation abrogates GSK-3-mediated phosphorylation and degradation of AJUBA. **A.** Western blot analysis comparing protein expression levels in wild type (WT) intestinal epithelial tissue, wildtype organoids, Apc LOF organoids, and Bcat GOF organoids. Two replicates shown for each experimental condition. β-Actin is used as a loading control. **B.** Immunofluorescence staining of organoids showing increased nuclear YAP localization (red) in Apc LOF relative to Bcat GOF organoids. Scale bar, 50 µm. **C.** Schematic of AJUBA protein showing reported phosphorylation sites (blue vertical lines), including serines-25, 39, 69, 79, 104, 119, 122, 123, 126, 127, 128, 133, 136, 137, 155, 175, 196, 202, 230, 237, 263 and threonines-30, 114, 162, 265. Magnified sequence shows multiple GSK-3 phosphorylation consensus motifs, illustrated by “SxxxSxxxS” where “S” indicates serine or threonine and “x” indicates any amino acid. Red text indicates known phosphorylated residues. Red * indicates sites identified in Shinde et al. 2017. **D.** Phosphorylation of AJUBA by recombinant GSK-3β using HEK293T cells ectopically expressing AJUBA. Immunoprecipitated AJUBA was added to an *in vitro* kinase reaction with γ-[32P]-ATP with or without recombinant GSK-3β. A representative of n = 3 biological replicates is shown. **E.** 293T cells were treated with the GSK-3 inhibitor CHIR99021 (9 µM) or vehicle control for 0, 24, or 48 hours and then immunoblotted for AJUBA and GAPDH. Analysis was performed in triplicate using LICOR and a representative immunoblot is shown. Quantification of replicate western blots is shown on the right panel. Intensity of AJUBA relative to GAPDH was normalized to time 0 value. **F-G.** SW480 cells were treated with cycloheximide (CHX) alone or with CHIR99021 for 0, 2, 6, 8, and 12 hours, and immunoblotted for AJUBA or β-Actin. Analysis was performed in triplicate and a representative immunoblot is shown (F), quantified in (G). Intensity of AJUBA relative to β-Actin was normalized to time 0. **H.** SW480 cells expressing control vector (YFP), APC SAMP motifs (SAMP), or full-length APC were treated with CHIR99021 or vehicle control as indicated, followed by immunoblotting 24 hours later. A representative immunoblot from 4 biological replicates is shown. **I.** Quantification of replicates from (H). The mean AJUBA protein relative to GAPDH is shown, normalized to control. n = 4 replicates. **J.** Western blot of freshly isolated intestinal epithelial lysates from wildtype, *Gsk-3a/b^flx/flx^::VillinCreER*, and *Apc^flx/flx^::VillinCreER* mice after tamoxifen injection, immunoblotted for AJUBA and GAPDH. **K.** Quantification of n=3 replicate Western blots from (J). AJUBA intensity relative to loading control was normalized to WT value. n = 3 wt, n = 3 GSK-3 KO, n = 5 APC KO. **L.** Clonal organoid-forming capacity of Apc LOF cells infected with *Ajuba* shRNA virus (sh-Ajuba) compared to control (ctrl) over three passages. **M.** UMAP plot showing single cell transcriptomes from indicated genotypes, including Apc LOF organoids (gray), Bcat GOF organoids transfected with AJUBA overexpressing (OE) virus (blue), or control virus (green), and shared secretory cells (orange). **N.** GSEA plot showing the enrichment of the fetal intestinal gene signature in Bcat GOF organoids overexpressing AJUBA relative to control Bcat GOF organoids as in (M).

We therefore asked whether established regulators of YAP may represent novel GSK-3 targets. AJUBA is a Lim domain-containing protein^52^ that suppresses LATS, which itself is a negative regulator of YAP activity, making AJUBA a positive regulator of YAP transcriptional activity^53–55^. AJUBA has multiple GSK-3 consensus motifs, including one identified in a prior phosphoproteomic study screening for novel GSK-3 targets in pluripotent cells^56^ **(Fig. 4C)**. Further, AJUBA protein was elevated relative to normal tissue in the vast majority (53/67) of CRC tumors examined in Jia et al^57^, where it was found to be important for survival in CRC cell lines.

We tested whether AJUBA is a direct substrate of GSK-3. *In vitro* phosphorylation assays with recombinant GSK-3β, AJUBA, and radiolabeled phosphate indicate robust AJUBA phosphorylation by GSK-3 (**Fig. 4D**). Pharmacological GSK-3 inhibition in HEK293 cells (*APC* wildtype with an intact WNT pathway) resulted in AJUBA accumulation, and was accompanied by increased AJUBA-LATS interaction, consistent with the published model of AJUBA sequestering LATS to inhibit its phosphorylation of YAP (**Fig. 4E** and **S5A**). To assess the effect of GSK-3 phosphorylation on AJUBA protein turnover, we inhibited translation and measured AJUBA protein over 12 hours. AJUBA levels declined with a half-life of ∼10 hours, and this was blocked by GSK-3 inhibition, suggesting that phosphorylation by GSK-3 accelerates AJUBA degradation (**Fig. 4F**, **G**). Next, colon cancer derived SW480 cells, harboring oncogenic APC mutations, were transfected with full-length *APC* and ^58–61^then treated with GSK-3 inhibitor. Restoring WT APC expression reduced AJUBA levels, and GSK inhibition potently blocked this, elevating AJUBA levels beyond control levels (**Fig. 4H**, **I**). AJUBA protein levels were also reduced by expression of a fragment APC protein containing the SAMP (Ser-Ala-Met-Pro) motifs, which is sufficient to activate GSK-3, restore β-CATENIN degradation, and rescue APC LOF phenotypes. Inhibition of GSK-3 similarly blocked the effect of the SAMP domain, elevating AJUBA protein levels above control. Taken together, these data indicate that APC potentiates GSK-3 activity to promote AJUBA turnover and suppress YAP activity whereas loss of APC or direct inhibition of GSK-3 stabilizes AJUBA. We verified this *in vivo* by inducing genetic ablation of either *Apc* or *Gsk-3a/Gsk-3b* throughout the intestinal epithelium using *Villin-CreER* and confirmed that loss of *Apc* or *Gsk-3a/b* stabilizes AJUBA protein (**Fig. 4J, K**).

We also asked whether the regulation of YAP by APC is evolutionarily conserved in vertebrates. Indeed, zebrafish larvae harboring a truncating mutation in *apc* (*apc^mcr^*^62,63^) exhibited increased Yap and Taz protein levels as well as their downstream target genes (**Fig. S5B-E**). Many of these Yap target genes^64^ were also found to be upregulated in APC LOF intestinal organoids and overlap with genes in the fetal intestinal signature (**Fig. S5F**).

Finally, we asked whether AJUBA contributes to β-CATENIN-independent phenotypes of APC LOF, and whether activation of AJUBA can induce fetal gene expression in β-CATENIN GOF organoids. Indeed, short hairpin RNA-mediated knockdown of AJUBA in APC LOF organoids suppressed clonal organoid forming capacity without altering proliferation (**Fig. 4L and S5G**). Conversely, forced expression of AJUBA in β-CATENIN GOF organoids was sufficient to drive expression of the fetal gene expression signature. (**Fig. 4M, N**). However, these organoids remained transcriptionally distinct from APC LOF organoids, supporting the notion that APC loss has additional β-CATENIN-independent effects beyond AJUBA-YAP stabilization (e.g., the previously described activation of mTORC1).

## DISCUSSION

The tumor suppressor *APC* is the fifth most commonly mutated genes across all cancers^65^, primarily due to the prevalence of colorectal cancer and its high rate of mutation (>80%) in CRC. For over two decades, numerous studies have focused on a single oncogenic consequence of APC inactivation, the aberrant stabilization of the canonical WNT pathway effector β-CATENIN. The field has largely overlooked the potential for β-CATENIN-independent oncogenic consequences of APC inactivation due to the fact that mouse models of β-CATENIN hyperactivation do exhibit hyperplasia and can drive tumorigenesis, and much attention has been given to the role of canonical WNT activity as a mitogen and self-renewal factor in the adult colonic stem cell compartment^63^. However, to date efforts at therapeutic intervention in APC-mutant CRC via targeting WNT^High^ carcinoma cells have been largely fruitless with tumors rapidly replacing ablated WNT^High^ cells from an otherwise WNT^Negative^ population^67^. Given the clinical observation that APC inactivation is the predominant initiating event in CRC, in contrast to activating mutations in *CTNNB1*, compounded with the fact that the destruction complex responsible for degradation of β-CATENIN in normal intestinal epithelium contains two kinases, GSK-3 and CK1α, which have scores of targets beyond β-CATENIN^68,69^, it is reasonable to hypothesize that APC loss may have oncogenic consequences beyond β-CATENIN stabilization and these effects may be mediated by dysregulation of GSK-3 and CK1α upon targets beyond β-CATENIN.

Here we employ mouse genetic and organoid models to demonstrate that APC inactivation is clearly more potent at promoting tumor initiation and progression in CRC models relative to hyperactivation of β-CATENIN. Unbiased transcriptomic analysis reveals that APC inactivation potently suppresses the pro-differentiation transcriptional regulator HNF4, and perhaps more strikingly, potently induces a fetal intestinal gene expression program associated with activity of the Hippo pathway transcriptional effector YAP, entirely independently of β-CATENIN transcriptional activity. YAP is an important oncogene in a host of cancers, including CRC, and its activation is associated with chemotherapy resistance, EMT-like phenotypes, and metastatic proclivity. Interestingly, YAP activation has previously been observed downstream of APC inactivation^15,70^. For example, Azzolin et al. showed that activation of Wnt signaling releases YAP/TAZ from the APC complex to enhance YAP/TAZ target gene expression^70^. Separately, Cai et al.^15^ showed that APC, acting through GSK-3, promotes LATS1-mediated phosphorylation and degradation of YAP. However, the direct substrate of GSK-3 in this regulation was not identified. Here we show that AJUBA, which interacts with and suppresses the function of LATS1^71^, is a direct target of GSK-3. Our data support a mechanism whereby APC potentiation of GSK-3 activity^12^ promotes the phosphorylation and degradation AJUBA^54,55^, providing a link between APC, LATS1, and the regulation of YAP protein abundance. Conversely, the loss of APC observed in CRC impairs GSK-3 activity, causing accumulation of AJUBA, reduced activity of LATS1, and increased YAP abundance. Consistent with both Azzolin et al.^70^ and Cai et al.^15^ elevated YAP then translocates to the nucleus to activate YAP-dependent fetal gene expression.

Phenotypically, the dual YAP-β-CATENIN activation we observe downstream of APC loss confers both an increase in proliferation relative to normal epithelium, known to be driven by the mitogenic activity of β-CATENIN, as well as enhanced stem cell and metastatic phenotypes associated with YAP activity. A number of recent studies have described the requirement for YAP activity for metastatic colonization and the converse requirement for β-CATENIN activity for proliferative outgrowth thereafter^67,72^. Further, the plasticity of fetal reversion conferred by YAP (along with SOX9 in the context of CRC) has been described in numerous studies (reviewed in ^16^). While a number of external signals have been implicated in inducing this fetal reversion, including fibroblast-derived prostaglandin E2 in the context of adenoma initiation^73^ and macrophage-derived TGFβ or T cell-derived IFN-γ in the context of post-injury regeneration^26,27^, our data demonstrate that APC inactivation is sufficient to induce YAP transcriptional activation and fetal plasticity in an epithelial cell-autonomous manner. It is, however, important to note that rare tumors driven by activating β-CATENIN mutations are still likely to contain cells in the YAP-driven fetal state due to the presence of these microenvironmental signals *in vivo*.

One striking finding in our current study was the remarkable ability of APC-null cells in the WNT/ β-CATENIN/adult stem cell-like state to move into the YAP/fetal-like stem cell state over the course of days. This finding may help to explain findings such as those of De Sousa e Melo et al. who observe that ablation of Lgr5+ WNT/ β-CATENIN cancer stem cells has little effect on the tumor and that this population is rapidly regenerated after the removal of the negative selective pressure on Lgr5+ cells^67^. Our findings support a model whereby APC loss generates a bistable epithelial population where cells fluctuate between the β-CATENIN/adult stem cell-like state and the YAP/fetal-like stem cell state, and this accounts for the propensity for colorectal adenocarcinomas to be initiated by mutations resulting in APC loss rather than β-CATENIN stabilization. Clinically, our findings support a therapeutic approach targeting both β-CATENIN and YAP for concomitant repression.

## ACKNOWLEDGEMENTS

We thank Dr. Eric Fearon (University of Michigan) for providing the *CDX2-G22-Cre* mouse strain and Dr. Enrico Radaelli and the comparative pathology core at the University of Pennsylvania School of Veterinary Medicine for blinded analyses of intestinal phenotypes in β-CATENIN and APC mouse models. The *Col1a^TetOdnTcf4^::R26^m2rtTA^* mouse strain was generated in conjunction with the University of Pennsylvania School of Veterinary Medicine Center for Germ Cell Research and Animal Transgenesis. This work was funded by R01CA168654 to CJL, R50CA221841 to NL, R01GM115517-01 to PSK, and the Harriet Ellison Woodward endowed chair (CJL). RLM was support by a T32 from the NIDDK (T32DK007780). This work was supported by core facilities, particularly the Molecular Pathology and Imaging Core associated with the NIDDK P30 Center in Molecular Studies in Digestive and Liver Diseases at the University of Pennsylvania Perelman School of Medicine (P30DK050306).

## METHODS

### Mouse Models

Generation of mice with doxycycline-inducible expression of N-terminally truncated TCF4 (dnTCF) protein (*Col1a^TetOdnTcf4^::R26^m2rtTA^*). To generate mice with dox-inducible expression dominant-negative TCF4 (dnTCF), we inserted a mouse N-terminally truncated *Tcf4* sequence under the control of the tetracycline operator with a minimal CMV promoter as described in^44,45^. This construct was targeted to the safe-haven chromatin downstream of the *Coll1a1* locus in mouse KH2 embryonic stem (ES) cells (129/sv × C57Bl/6 F1 hybrid male) harboring a modified reverse tetracycline transactivator (M2rtTA) targeted to and driven by the endogenous *Rosa26* locus. The insert sequence, acquired from the EdTC plasmid (Roel Nusse, Fuerer et al.), was cloned into the pBS31’-RBGpA backbone using In-Fusion cloning. The first 31 amino acids of the N-terminus were deleted, resulting in a dominant-negative form of TCF4 that lacks the β-Catenin binding domain, analogous to that described in ^33^. Protein expression was verified by dox administration (2ug/mL). The resulting *Col1a^TetOdnTcf4^::R26^m2rtTA^* were then injected into C57Bl/6 blastocysts, and resulting chimeras were the backcrossed to C57Bl/6 mice to generate the *Col1a^TetOdnTcf4^::R26^m2rtTA^* mouse strain from which intestinal organoids were derived. The *Apc^flox^* allele is described in^28^, the *Ctnnb1^ex3flox^* allele is described in^29^, the *Villin-CreER* allele is described in^31^, the *CDX2-G22-Cre* allele is described in ^35^, the GSK-3α/β floxed alleles are described in ^74^.

### Mouse Crypt Isolation and Organoid Culture

Murine intestinal organoids were cultured following^75^, with a summary provided here. *Apc^flox/flox^::VillinCreER* and *Ctnnb1^ex3flox/ex3flox^::VillinCreER* mice were administered 100 µl of Tamoxifen (10 mg/ml, Millipore Sigma, T5648) daily for three consecutive days prior to crypt collection. Age- and sex-matched wild-type mice from the same C57BL/6J background (The Jackson Laboratory, stock no: 000664) were used as controls. Crypts were isolated from the first 2–12 cm of the intestine or distal colon by incubating the tissue with 5 mM EDTA (Invitrogen, 15575-020) in Hank’s balanced salt solution at 4 °C for 15 minutes with gentle rotation (12 cycles/min). After incubation, crypts were released by gently passing the tissue through the narrow gap between a serological pipette and the bottom of a 50 ml Falcon conical tube. The crypts were then washed twice with 0.5% BSA in HBSS and collected by slow centrifugation at 800 rpm for 5 minutes at 4 °C. Approximately 500 crypts were mixed with 100 µl Matrigel and seeded into one well of a 24-well plate. After allowing the Matrigel to solidify in an incubator for 10 minutes, each well received 200 µl of media. Organoid media includes advanced Dulbecco’s modified Eagle medium/F12 (Thermo Fisher Scientific, 12634-010) supplemented with N2 (Invitrogen, 17502-048), B27 (Invitrogen, 17504-044), N-acetylcysteine (MilliporeSigma, A9165, 1 mM), mEGF (Invitrogen, PMG8041, 50 ng/ml) and 50% L-WRN (ATCC, CRL-3276) conditional medium. In instances where cultures are derived from single cells, Jagged-1 (AnaSpec, 61298, 1 mM), Y-27632 (Selleckchem, S1049, 10 mM), and CHIR99021 (Stemgent, 04-0004, 5 mM) are added for the first 48 hours only.

### EdU Assay

For S-phase analysis, organoids/tumoroids were digested into single cells using TrypLE and seeded into a 24-well plate at a density of 5,000 cells per well, with each cell type seeded in triplicate. Organoids were cultured for four days, with media changed daily. On day four, 10 µM EdU (Thermo Fisher #E10187) was added to the culture media. Cells were harvested two hours after EdU incorporation. Single-cell suspensions were prepared using TrypLE digestion for flow cytometric analysis. EdU staining was performed using the Click-iT Plus EdU Alexa Fluor 647 Flow Cytometry Assay Kit (Thermo Fisher #10634) following the manufacturer’s protocol. Stained cells were analyzed on an LSR Fortessa (BD Biosciences), and data analysis was performed using FlowJo software (BD Biosciences).

### FACS Sorting and 10x Genomics Single-Cell RNA-seq (scRNA-seq) Library Preparation

Cultured organoids were digested with TrypLE™ Express Enzyme (ThermoFisher, 12605010) for 5-8 minutes at 37°C. The cell pellet was examined under a microscope to verify quality before sorting. Cells were resuspended in HBSS with 0.5% BSA and sorted using a Becton Dickinson (BD) FACS Aria II controlled by BD FACS DIVA software. FSC-H, FSC-W, SSC-H, and SSC-W parameters were used in combination to exclude doublets. Sorted cells were collected into a protein low-binding tube (Eppendorf, 0030108442) containing HBSS with 0.04% BSA. Cells were immediately counted using an Invitrogen Countess™ II Automated Cell Counter to confirm quality and then loaded into the 10x Genomics Chromium following the manufacturer’s instructions (Chromium Single Cell 3’ Reagent Kits v3.1). Multiple samples within each library were multiplexed using cell hashing (BioLegend, TotalSeq™-B) or 10x Chromium Cell Multiplexing Oligos. The 3’ Gene Expression and Cell Multiplexing libraries were pooled together for sequencing at a 4:1 ratio. Sequencing was performed on an Illumina NovaSeq 6000, aiming for a minimum coverage of 50,000 reads per cell (paired-end; read 1: 28 cycles; i7 index: 10 cycles; i5 index: 10 cycles; read 2: 90 cycles).

### Immunofluorescence, Whole Mount Staining, and Microscopy

For 2D imaging, tissue samples and organoids were fixed in 4% paraformaldehyde aqueous solution (Electron Microscopy Sciences) overnight (14–16 hours), followed by alcohol dehydration, xylene clearing, and paraffin embedding. Organoid samples were initially embedded in 4% low melting temperature agarose (ThermoFisher) to maintain their shape before paraffin treatment. Paraffin blocks were cut into 5 µm sections, adhered to glass slides, heated at 70°C for 1 hour, deparaffinized in xylene three times, and rehydrated in a graded alcohol series (100%, 90%, 80%, 70%). Antigen retrieval was performed using pH 9.0 Tris-EDTA buffer boiled in a pressure cooker for 3 minutes. Slides were blocked at room temperature for 1 hour with 10% normal serum and permeabilized with 0.5% Triton X-100 in Tris-buffered saline. Primary antibodies were incubated overnight at 4°C. Following TBST washes, sections were incubated with secondary antibodies (Cy2-, Cy3-, and Cy5-conjugated; Jackson Immunoresearch Laboratories), stained with DAPI, and embedded using VECTASHIELD (Vector Labs). Raw images were captured on a Leica DMi8 microscope equipped with DFC9000GT and MC170 HD cameras and processed using Leica Application Suite X (LAS X) or ImageJ software.

### Endoscope-Guided Orthotopic Transplantation

Approximately 8-week-old C57BL/6J mice from the Jackson Laboratory were used for orthotopic transplantation experiments. Organoids derived from *Apc^flox/flox^*::*Villin-CreER* (APC LOF) and *Ctnnb1^ex3flox/ex3flox^*::*Villin-CreER* (β-CATENIN GOF) mice, with or without *Tp53* inactivating mutations^76^, were recovered from Matrigel using Cell Recovery Solution (Corning #354253) by incubating on ice for 30 minutes. Cell clusters, equivalent to 2 × 10^4 cells per mouse, were resuspended in cold DPBS with 0.5% BSA and 10% Matrigel. These were then transplanted into the colon mucosa of recipient mice via optical colonoscopy using a custom injection needle (Hamilton Inc., 33 gauge, small Hub RN NDL, 12 inches long, point 4, 45-degree bevel, catalog #7803-05), a 100 µl syringe (Hamilton Inc., part number 7656-01), a transfer needle (Hamilton Inc., part number 7770-02), and a colonoscope with an integrated working channel (Richard Wolf 1.9 mm/9.5 French pediatric urethroscope, part number 8626.431). A 70 µl cell suspension was delivered to the colon mucosa, with one injection performed per mouse. Mice underwent colonoscopy 2-8 weeks following organoid transplantation to assess tumor development. Tumor size was quantified in situ by photographing the tumors and measuring the percentage of the lumen occluded by the tumor. At the endpoint, mice were sacrificed, and tumor weights were recorded. Liver metastasis was also evaluated after dissection.

### Slide Evaluation

All slides were reviewed by three board-certified veterinary pathologists, blinded to condition, and the results represent the diagnoses reached by consensus. MMHCC references were used for the classification of mouse proliferative lesions in the GI tract. The main microscopic changes observed in the set of slides provided are characterized by aberrant hyperplasia involving various compartments along the crypt-villus unit of the small intestine. The following semiquantitative scoring system has been developed and applied to assess nature, distribution, and severity of these chances across the different samples. Combined scores are reported in the Results.

Scoring System

extension affected:

- - no or sporadic evidence of hyperplasia/dysplasia in the compartment.
- - 5 to 20% of the compartment affected by hyperplasia/dysplasia.
- - 21 to 50% of the compartment affected by hyperplasia/dysplasia.
- - more than 50% of the compartment affected by hyperplasia dysplasia.

mitotic index (average number based on the evaluation of 10 affected compartments):

- - mitosis ≤1.
- - mitosis >1 and <3.
- - mitosis ≥3 and <6.
- - mitosis ≥6

degree of hyperplasia

- - no evidence of hyperplasia
- - mild hyperplasia characterized by a regular expansion/elongation of the affected compartment.
- - moderate atypical hyperplasia characterized by irregular expansion/elongation of the affected compartment but overall preservation of the normal epithelial architecture.
- - marked atypical hyperplasia/dysplasia characterized by irregular expansion/elongation of the affected compartment with extensive loss of the normal epithelial architecture.

### Western blot analysis

Organoids were treated with Cell Recovery Solution, washed with D-PBS, pelleted, and lysed with 1x RIPA Buffer (CST#9806) containing a protease/phosphatase inhibitor cocktail (CST#5872S). Organoids were disrupted by sonication and centrifuged for 15 minutes at 14,000 g. The supernatant was transferred to a clean tube, and protein concentration was determined using a BCA assay (Thermo Fisher Scientific). An appropriate volume of 4x loading buffer was added to the cell lysates, followed by incubation at 98°C for 10 minutes. Equal amounts of protein were loaded into each lane of an 8–12% SDS–PAGE gel. Membranes were blocked with 5% milk for 1 hour at room temperature, then incubated overnight at 4°C with the indicated primary antibodies diluted in 5% milk. This was followed by incubation with HRP-conjugated secondary antibodies for 1 hour. Western blotting analyses were performed using the following primary antibodies: Phospho-YAP/TAZ Antibody Sampler Kit (CST#52420T), Yap (D8H1X) XP® Rabbit mAb (CST#14074S), ZMYND8 Polyclonal Antibody (Thermo Fisher Scientific#11633-1-AP), Ajuba (CST#4897), Flash Phalloidin Green 488 (BioLegend#424201) and Hippo Pathway: Upstream Signaling Antibody Sampler Kit (CST#56612), which includes antibodies against TAZ (D3I6D) Rabbit mAb (CST#70148), Ajuba (D4D8P) Rabbit mAb (CST#34648), and Anti-rabbit IgG, HRP-linked antibody (CST#7074). Signals were detected with HRP-conjugated secondary anti-rabbit antibody (CST#7074S, 1:2,000) and anti-mouse antibody (CST#7076S, 1:5,000) and visualized using the SuperSignal West Pico PLUS Chemiluminescent Substrate (Thermo Scientific#34577) and the Bio-RAD Chemidoc TMMP imaging system.

HEK293T Cells, HT-29-APC, HT-29-bgal, 293-STF, and SW480 cells were cultured in Dulbecco’s modified Eagle’s medium supplemented with 10% fetal bovine serum and 1% pencillin/streptomycin. Cells were transfected 24hr after plating using Lipofectamine 3000 for plasmids. Where indicated, cells were treated with 50 µM cycloheximide (CHX) and/or 9 µM CHIR99021. Cells were lysed in buffer containing 20mM Tris pH 7.5, 140mM NaCl, 1mM EDTA, 10% glycerol, 1% Triton X-100, 50mM NaF, 1mM DTT, protease inhibitor cocktail, phosphatase inhibitor cocktail. Supernatants were collected after centrifugation at 13,000rpm for 15min at 4 °C. Samples were heated at 100 °C for 5min before SDS-Page and Western Blot Analysis. Western blots were imaged with the LICOR Odyssey system and quantified using Image Studio software.

Zebrafish embryos were raised at 28.5°C in standard E3 medium. The chorion was manually removed at 24 hours post-fertilization (hpf). Rapamycin was added directly to the medium starting at 24 hpf. Medium was changed and fresh treatments were added every 24hours. Zebrafish husbandry and egg procurement were carried out in accordance with the guidelines of the University of Pennsylvania Institutional Animal Care and Use Committee. Zebrafish embryos were lysed on ice in 5 µl/embryo of buffer containing 1% NP-40, 20mM Tris pH 8.0, 50mM NaCl, 2.5mM EDTA, 1mM DTT, protease inhibitor cocktail, and phosphatase inhibitor cocktail. 2X Laemmli sample buffer was added, samples were heated at 100 °C for 5min, and centrifuged at 14,000rpm for 5min at 4 °C. Supernatants were collected and 1 embryo equivalent (∼10µl) was loaded per lane for SDS-PAGE and western blot analysis as above.

### *In vitro* kinase assays

Purified recombinant GSK-3 (from Cell Signaling Technologies) and AJUBA (immunoprecipitated from HEK293T cells transfected with AJUBA expression plasmid) were incubated in protein kinase reaction buffer (100mM Tris pH 7.5, 5mM DTT, 10mM MgCl2, 100µM ATP) for 30 min at 30°C. Reactions were stopped by adding 4X Laemmli Sample Buffer. Samples were resolved on SDS-PAGE and gels were fixed directly in 10% acetic acid and 10% methanol overnight, dried, and exposed to X-ray film.

### Bulk RNA seq

Freshly isolated intestinal crypts from *Apc^flox/flox^::VillinCreER* and *Ctnnb1^ex3flox/ex3flox^::VillinCreER* and control wild-type mice were dissolved in TRIzol (Thermo Fisher Scientific). Following phase separation, the aqueous phase containing RNA was mixed with 70% ethanol and transferred to RNeasy MinElute spin columns (QIAGEN) for RNA cleanup and concentration using the RNeasy MinElute Cleanup Kit (QIAGEN). cDNA synthesis was performed using the SMART-Seq v4 Ultra Low Input RNA Kit for Sequencing (Clontech). Libraries were prepared with the Nextera XT DNA Library Prep kit (Illumina) and sequenced on an Illumina NextSeq 550 instrument, generating 20-40 million high-quality 75-bp single-end reads per sample. Adapters were trimmed using Trimmomatic v0.32^77^ and reads were aligned to the mm10 genome using STAR v2.3.0e^78^. Samtools v0.1.19^79^ was used for file conversion, alignment filtering, sorting, and indexing. Read quantification per gene was performed using Rsubread’s feature counts command^80^ with Ensembl mm10 annotations, removing genes with CPM ≤ 1 in at least a quarter of the samples. Data were log2 transformed and normalized using Limma v3.48.3’s voom function^81^ and differential expression analysis was conducted with Limma. Pathway enrichment was analyzed using Camera^82^ with EnsDb.Mmusculus.v75 via RNA-webInterface (https://github.com/bapoorva/RNA-WebInterface). Sample variability was visualized using Principal Component Analysis (PCA). In an alternative analysis, raw data were mapped to the mouse reference transcriptome using Kallisto^83^ and analyzed with RUVseqr^84^ and edgeR^85^ in R. Pathway enrichment was determined using GSEA with the MSigDB Collection 2^37^. RNA-sequencing data have been deposited in the NCBI Gene Expression Omnibus (GEO)^86^, accession number GSE266579. RNA-seq on WT and *apc^mcr/mcr^* zebrafish larvae (3 days post-fertilization) was described previously^87^. RNA-seq data were deposited in the NCBI Gene Expression Omnibus and are accessible through GEO Series accession number GSE234986.

### Single Cell RNAseq Data Processing

Sequencing reads for samples were initially pre-processed using the 10x Genomics Cell Ranger pipeline. Reads were aligned to the respective reference genomes, GRCh38 for human and GRCm38 (mm10) for mouse. The raw gene-barcode matrix produced by the Cell Ranger ‘cellranger count’ function underwent initial filtering to remove barcodes with fewer than 1000 transcripts (quantified by unique molecular identifiers (UMIs)) and fewer than 500 expressed genes (where “expressed” indicates at least one transcript from the gene is present in the cell). Barcodes passing this filter were considered as cells and included in the final dataset. Additionally, cells exhibiting high levels of mitochondrial gene expression (exceeding 20% of the total UMI count) were excluded from all subsequent analyses. For samples multiplexed using the TotalSeq-B protocol, cells were demultiplexed by performing Louvain clustering on the UMAP generated from the hashtag count matrix. The gene-barcode UMI count matrices from all datasets were then size-factor corrected and log-transformed to generate a normalized gene expression matrix.

We used the Seurat^88^ and VisCello^89^ packages to generate a series of principal component analyses (PCAs) and uniform manifold approximation and projections (UMAPs) for different cell subsets. The processing pipeline followed the method described previously^90^. Briefly, we applied an “informative feature (IFF) selection” procedure to select genes with high Gini coefficients, indicating the inequality and specificity of gene expression across clusters. Principal component analysis (PCA) was then performed on the IFF-cell matrix, and the top principal components (PCs) were used as features for the UMAP algorithm. UMAP was computed using the ‘umap’ function in the ‘uwot’ R package, with a “cosine” distance metric, 30 nearest neighbors, and default parameters for the rest. Louvain clustering was performed on the k-nearest neighbor graph (k = 20) constructed from the cell embeddings in the UMAP. Each cluster was annotated by comparing cluster-specific differentially expressed genes with known cell-type marker genes from previous literature. Differential expression analysis was carried out using the “sSeq” algorithm, with FDRL<L0.05 and log2 fold change > 1 as cutoff for differentially expressed genes (DEGs).

### GSEA Analysis

Gene Set Enrichment Analysis (GSEA, Broad Institute)^37^ was used to determine whether defined sets of genes exhibit statistically significant, concordant differences between two biological groups (e.g., WT vs. Apc, WT vs. Ctnnb1, or Apc vs. Ctnnb1). Prior to GSEA, differential expression analysis was conducted using DESeq2 for bulk RNA-Seq data and Viscello for single-cell RNA-Seq data. For DESeq2, raw counts were normalized, and differentially expressed genes were identified based on fold change and adjusted p-values. For single-cell RNA-Seq, Viscello normalized the data and performed differential expression analysis considering the unique characteristics of single-cell data. Genes were then ranked according to their expression levels, and the ranked lists were input into GSEA for pre-ranked analysis mode. This assessed whether gene set members were randomly distributed throughout the ranked list or primarily found at the top or bottom, indicating enrichment. Enrichment scores for each gene set were calculated and compared to a null distribution generated through permutation testing to evaluate significance. Results were visualized using GSEA software to highlight significantly enriched pathways and biological processes, providing insights into the molecular mechanisms and pathways underlying the observed phenotypic differences.

### Fetal TCGA survival analysis

Patients from the COAD/READ TCGA dataset were scored according to their match to the fetal signature using the Singscore (1.26.0) R (4.3.0) package on default settings. The fetal upregulated and downregulated signatures were both used to generate a score representing the magnitude of match to the fetal signature. Scored TCGA patients were then divided into *APC* wt vs. mutant for further analyses. For both *APC* wt and mutant patients, Kaplan-Meier plots were generated showing overall survival trends in patients stratified into either (1) upper quartile vs. lower three quartiles match to fetal score, or (2) lower quartile vs. upper three quartiles match to fetal score. Log rank test were used to evaluate the significance of the differences using the Survival (3.8-3) R package.

### Data availability

All raw and processed sequencing data generated and analyzed in this study have been deposited in the Gene Expression Omnibus (GEO) database of the National Institutes of Health (NIH) under accession numbers GSE266579 and GSE234986. This ensures public availability, allowing other researchers to replicate and verify the results reported in this study. Human single-cell data compatible with VisCello has also been deposited to Zenodo46 (https://zenodo.org/record/7872684). Readers can use VisCello (https://github.com/qinzhu/VisCello) to interactively explore and analyze the deposited data, which include: an R data object (eset.rds) containing the raw count matrix, normalized expression matrix, and metadata for all cells; and the clist.rds file containing a list of dimension reduction results for different data subsets. The mouse transcriptome sequencing datasets generated are available in the NCBI Gene Expression Omnibus (GEO) under accession number GSE198759. Source data are provided with this paper.

The authors declare no competing interests.

---

**Figure S1.**
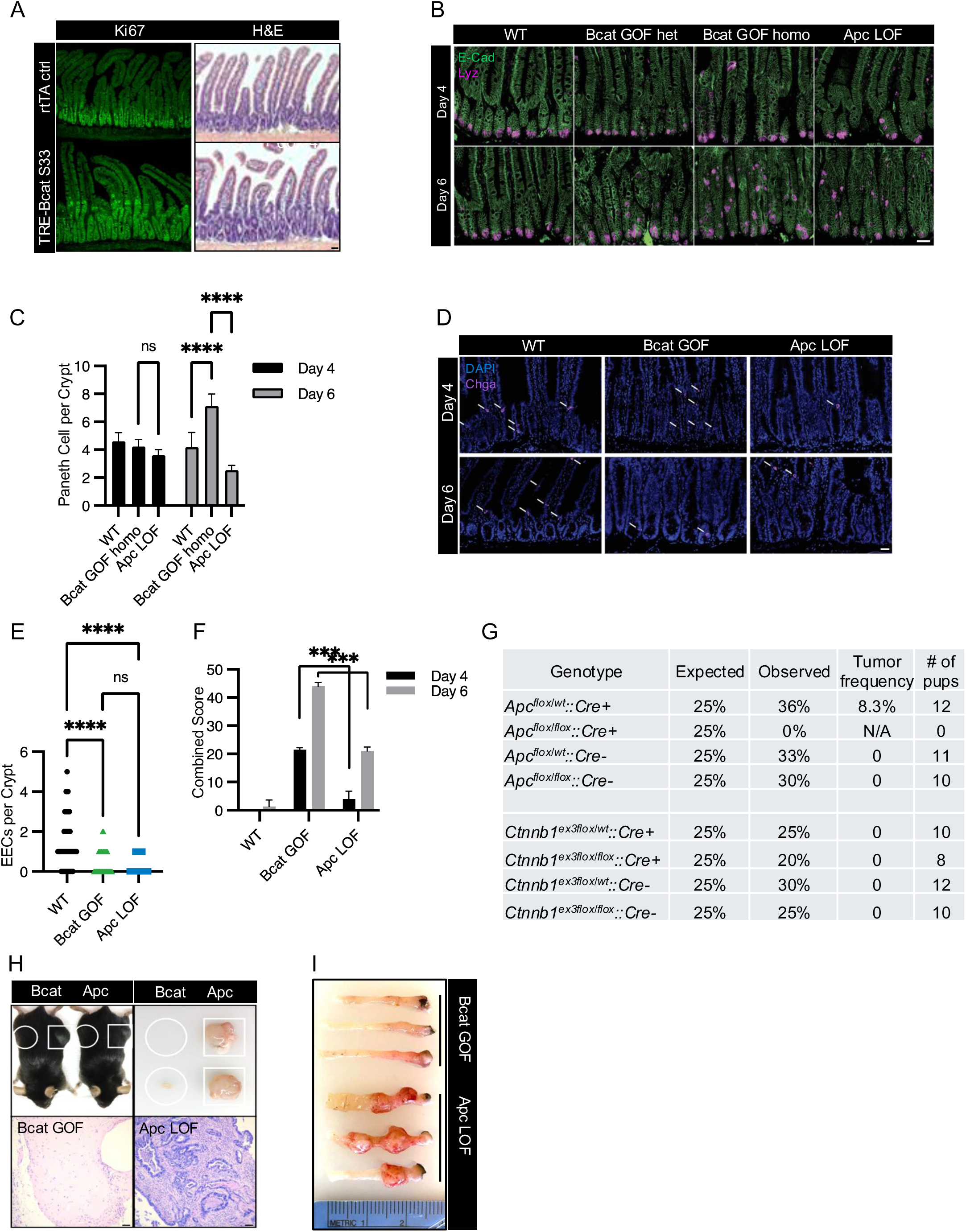
**A.** Histological sections of jejunum from *TRE-β-CATENIN-S33* mice after 7 days doxycycline induction, with proliferation visualized by KI67 immunofluorescence. Scale bar, 50 µm. **B.** Paneth cell stain in Apc LOF and Bcat GOF intestine of the indicated genotype at indicated timepoints post-tamoxifen induction. Scale bar, 50 µm. **C.** Quantification of Paneth cells per crypt shown in B. **D.** Immunofluorescence stain for ChgA+ enteroendocrine cells (EECs) in the jejunum of Apc LOF and Bcat GOF mice. Scale bar, 50 µm. **E**. Quantification of EECs per crypt shown in D. **F.** Combined histopathology scoring of affected area, mitotic index, and degree of hyperplasia in wild type (WT), Bcat GOF, and Apc LOF mice, performed by veterinary pathologists. **G.** Expected and observed birth frequencies of pups with different genetic combinations and visible tumor formation. **H.** Tumors resulting from subcutaneous implantation of Bcat GOF and Apc LOF organoids in recipient 2.5 months old mice. Scale bar, 50 µm. **I.** Representative images of tumors resulting from endoscopic orthotopic implantation of Bcat GOF and Apc LOF organoids into colonic mucosa 8 weeks post implantation. ****P < 0.0001, ***P < 0.001.

**Figure S2.**
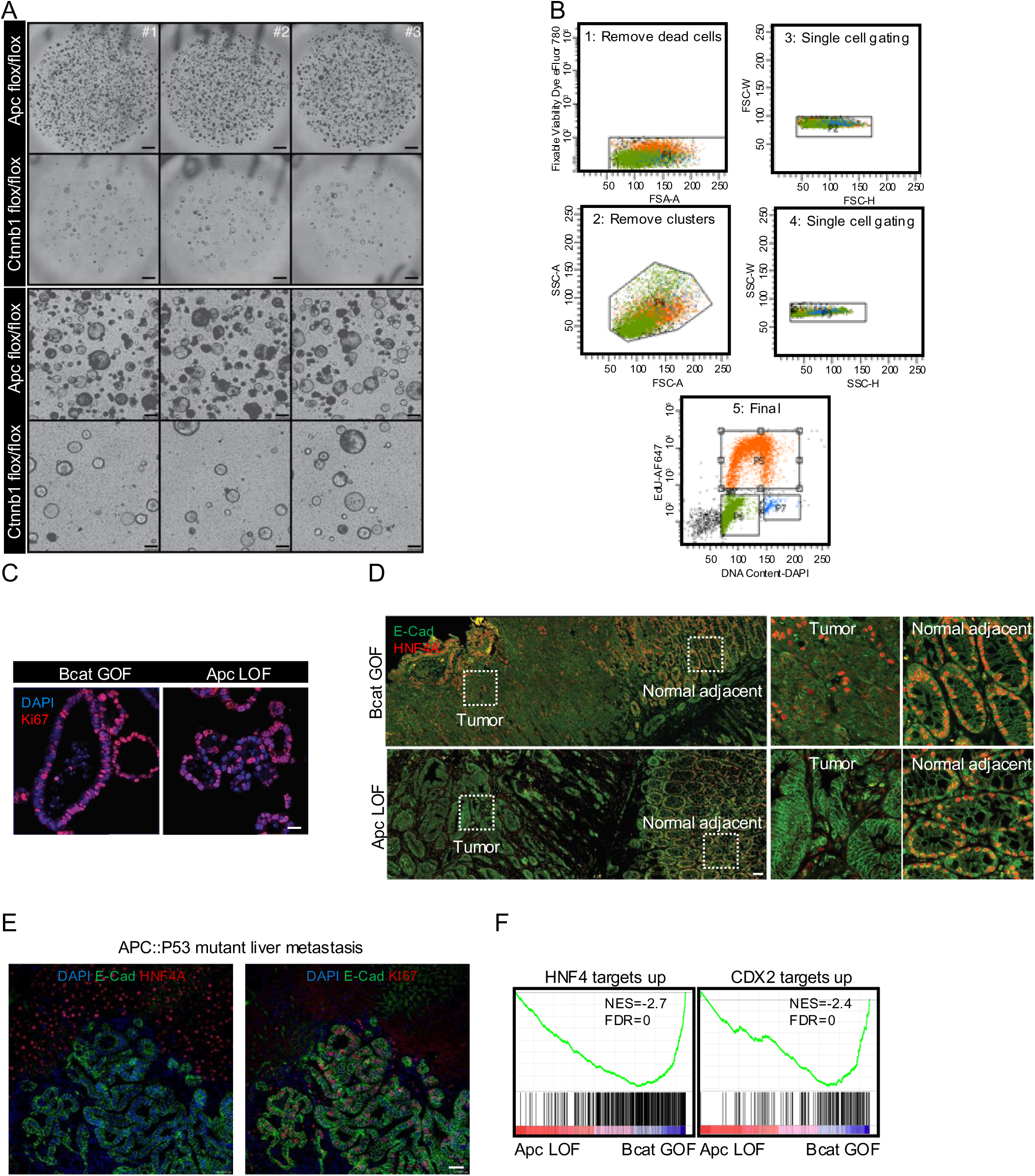
**A.** Representative brightfield micrographs of Apc LOF and Bcat GOF organoid cultures. Scale bar= 1mm (top panel) and 50 µm (bottom panel). **B.** Scatter plots delineating the stepwise gating strategy for flow cytometric analyses of organoid cultures. **C.** KI67 staining of representative Apc LOF and Bcat GOF organoids Scale bar= 50 µm. **D.** HNF4A and E-Cadherin immunofluorescence staining of tumors driven by Apc LOF or Bcat GOF and normal adjacent tissue, with inset zoom on right. Scale bar= 50 µm. **E.** HNF4A and E-Cadherin immunofluorescence staining in liver metastasis from primary tumors generated by orthoptic implantation of APC::P53 tumor organoids in the colonic mucosa. Scale bar= 50 µm**. F.** Gene set enrichment analysis (GSEA) comparing Apc LOF to Bcat GOF against gene sets representing positively affected target genes of HNF4A/G and CDX2. HNF4A/G targets are verified direct targets via chromatin immunoprecipitation.

**Figure S3.**
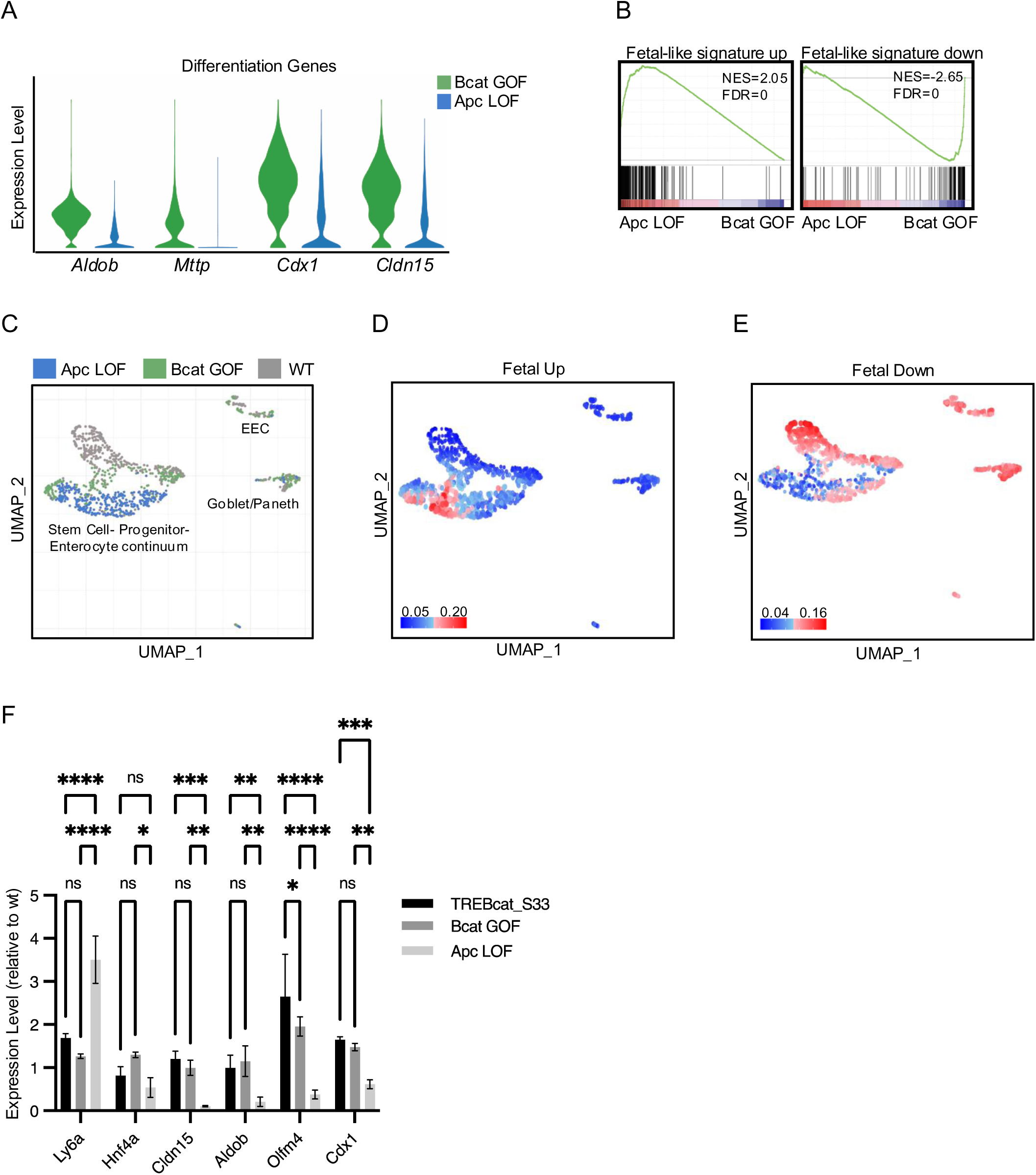
**A.** Violin plots showing expression of hallmark epithelial enterocyte differentiation genes derived from single cell transcriptomic analysis of Apc LOF and Bcat GOF organoid cultures. **B.** GSEA analysis examining enrichment of fetal signature (fetal signature up) and adult signature (fetal signature down) gene sets in Apc LOF and Bcat GOF organoid cultures. **C.** UMAP of single cell transcriptomes in Apc LOF, Bcat GOF, and wildtype control organoid cultures. **D, E.** UMAP plots as in C showing cell type distribution showing enrichment for fetal (fetal up, D) and adult (fetal down, E) intestinal signatures. **F.** Quantitative RT-PCR in Apc LOF, Bcat GOF, and Tre-BcatS33 in epithelial cells derived from mouse intestine showing expression relative to wildtype. Genes include hallmark fetal (*Ly6a*), differentiation (*Hnf4a, Cldn15, Aldob, Cdx1*), and Notch target/adult stem cell *Olfm4.* ****P < 0.0001, ***P < 0.001 **P < 0.01 *P < 0.05.

**Figure S4.**
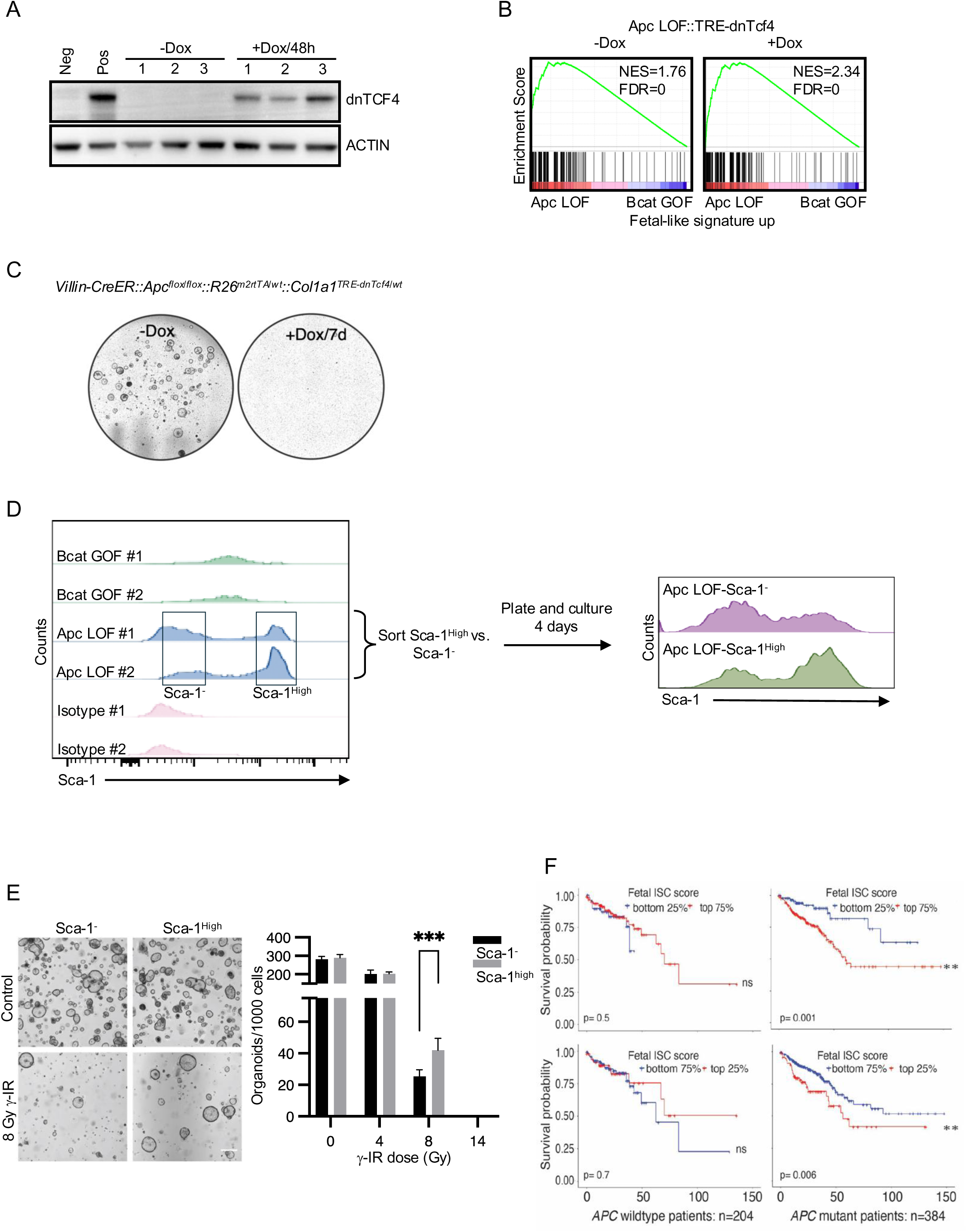
**A.** Western blotting for expression of dominant-negative TCF4 in targeted mouse embryonic stem cells in response to doxycycline (Dox) administration, showing three independently-targeted clones. **B.** GSEA for the fetal up gene signature in Apc LOF::TRE-dnTcf4 organoid cultures, with (right) or without (left) 48hrs Dox treatment, relative to Bcat GOF cultures. **C.** Organoid cultures from Apc LOF::TRE-dnTcf4 mice, with or without one week of Dox treatment. **D.** Histograms based on flow cytometric analysis of Sca1 expression in two replicates each of Bcat GOF cultures, Apc LOF cultures, and controls (left). Sca1^high^ and Sca1^-^ (negative) cells were then sorted, replated, cultured for 4 days and flow cytometric analysis repeated (right). **E.** Apc LOF organoids were broken down to single cells followed by sorting of Sca1^high^ and Sca1^-^ populations, either exposed to increasing doses of ionizing radiation (micrograph shown for 8 Gy exposure) or not (Control), then plated and cultured for 7 days. Clonal organoid efficiencies quantified at right. Scale bar, 50 µm. ***P < 0.001. **F.** Kaplan–Meier survival analysis of human patients, stratified by *APC* mutation status and Fetal ISC score. Survival probability is shown for *APC* wild-type (n=204) and *APC* mutant (n=384) patients, with comparisons between top and bottom quartiles of the Fetal ISC score. **P < 0.01, ns: not significant.

**Figure S5.**
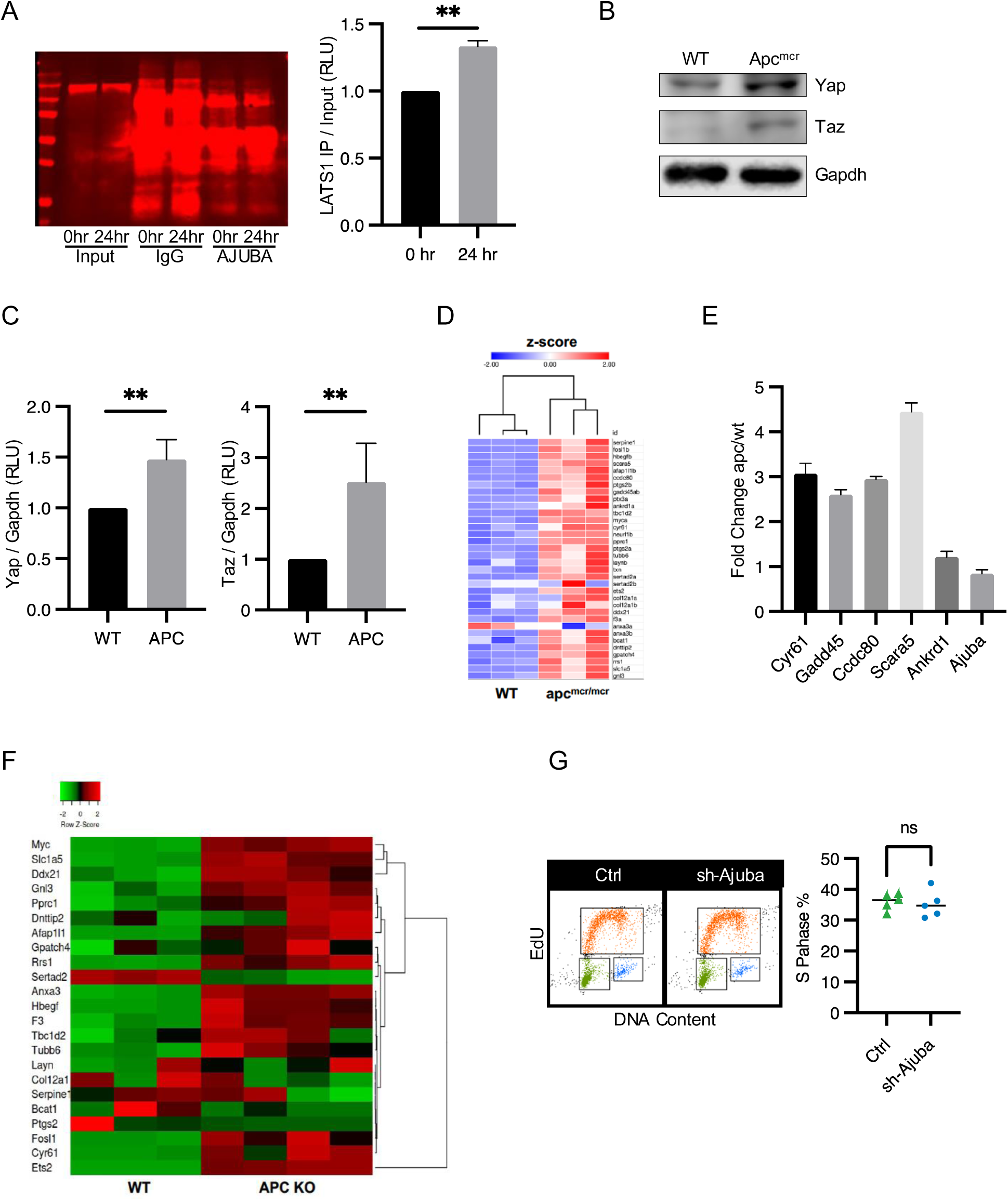
**A.** HEK293T cells were treated with 9 µM CHIR99021 for 0 or 24 hours, lysed, and incubated with anti-AJUBA antibodies or IgG control and immunoprecipitated then blotted for LATS1, quantified at right. **B.** 48 hpf wildtype and *Apc^mcr/mcr^*zebrafish were lysed and blotted =for Yap, Taz, and Gapdh. **C.** Quantification of blots from (B). **D.** Bulk transcriptomic analysis of wildtype and *Apc^mcr/mcr^*zebrafish at 72 hpf. Genes previously identified as YAP-TEAD or TAZ-TEAD target genes were identified, and fold change between wildtype and *Apc^mcr/mcr^*zebrafish larvae is shown. **E.** Expression of Ajuba and downstream Hippo target gene mRNA in ZF comparing wildtype to *Apc^mcr/mcr^* mutants. **F.** Bulk transcriptome profiling of wildtype and APC KO mouse intestinal epithelium for downstream Hippo target genes. G. FACS scatter plots for EdU assay in Apc LOF organoids infected with AJUBA shRNA of control virus.

